# Step-by-step methodology for building a capsule to sample small intestine content

**DOI:** 10.1101/2024.10.10.617430

**Authors:** Catherine Ollagnier, Marco Tretola, Inés García Viñado, Benoit Morel

**Affiliations:** Pig Research Unit, Agroscope, 1725 Posieux, Switzerland; iMedging, 1796 Courgevaux, Switzerland

**Keywords:** CapSa capsule, gut microbiome, ingestible sampling capsule

## Abstract

**Background:** Research on the intestinal microbiome has been hindered by limited access to intestinal content. Recently, a few capsule prototypes have demonstrated their potential for sampling intestinal material while using the natural pathway. However, access to these capsules is restricted because most of them are not yet commercially available. Pigs offer significant potential to inform human research due to the many physiological similarities between the two species. The unique features of pig anatomy have made it difficult to conduct research using swallowable devices. This article provides a detailed account of the manufacturing process and composition of a capsule, along with all the necessary steps for successfully sampling small intestine content in pigs.

**Results:** The capsule moves passively through the digestive tract, relying solely on intestinal peristalsis for propulsion. Engineered to open when it encounters a pH level greater than 6, the upper part dissolves, allowing intestinal fluids to enter the inner chamber. This triggers a plunger to expand, drawing luminal content into the storage chamber. Once the plunger mechanism is fully extended the capsule is automatically sealed. The capsule has a size of a 0 hard capsule and consists of two main components: a dissolvable exterior with an enteric coating and a 3D-printed bottom part. The printing files of the 3D-printed bottom part are provided for replication. In vitro testing shows that the capsule can withstand two hours in an acidic medium and successfully samples within an hour of being transferred to a neutral medium. When tested in vivo in pigs, the capsule successfully collected intestinal content from the upper and middle sections of the small intestine.

**Conclusions:** This article provides essential details for the rapid development of a cost-effective tool that has been already validated for non-invasive sampling of small intestine content in pigs. By providing access to the exact production steps and printing files, this article empowers others to innovate and expand upon this foundational work. This open-source approach opens up new avenues for intestinal research, making it more accessible and adaptable for a wide range of studies in both animal and human models.

## I. Background

The gastrointestinal tract (GIT) is not only the largest interface between the external and internal environments of animals, but it also contains the largest amount, and the greatest diversity of microorganisms [1]. The GIT microbiota is defined as an ecological community made up of commensal, symbiotic and pathogenic microorganisms, including bacteria, viruses, parasites, fungi, archaea and protists, that inhabit the mammalian gut [2]. Co-evolution of gut microorganisms with their hosts has led to the acquisition of microbial roles in digestion, nutrient utilization, toxin removal, protection against pathogens and regulation of the endocrine and immune systems [3, 4].

An important but often overlooked aspect of the gut is its regional heterogeneity, which not only impacts the local physiology but also results in very different microbial communities [5]. Because of difficulties in accessing and sampling the intestinal tract, stool has been the main source of information for human gut microbiome studies [6]. Indeed, the current gold standard for collecting a sample of gut microbiota is faecal sampling but it is not possible to associate a particular disease to a certain location within the length and the complexity of the GIT [7].

An immense potential has been described in the pig husbandry sector for manipulating the GIT microbiota to improve nutritional and immunological activities within the host to boost productivity [1]. The study of pigs has also a great potential to inform human research due to the many similarities identified in the physiological attributes of these two species. Pigs exhibit similar susceptibilities and clinical manifestations to pathogens that are the etiological agents in certain human intestinal disorders [8, 9].

The microbiota contains a depth of information that cannot be fully captured by fecal analysis, so capsules are being developed to collect samples from the gut to allow detailed examination. Devices that can collect a sample from the gut are divided into two broad categories. First, controlled or active sampling devices trigger the sampling process through wireless control (electronic and/or magnetic) to collect a sample at the target-site [10-13].

Second, uncontrolled or passive sampling devices activate the sample collection based on the physicochemical properties of their local environment.

The passive device capsules are intended to collect content from the lumen of intestines for ex vivo analysis. These capsules function primarily through an enteric coating that dissolves, triggering the sampling process at the target site. The characteristics of the enteric coating, particularly the time it takes to dissolve after reaching a specific pH, determine whether the capsule will sample fluid from the upper, middle, or lower section of the intestine. However, because the capsules rely on peristalsis to move through the gastrointestinal tract, the precise sampling location cannot be guaranteed. Once the outer membrane dissolves, the capsule’s aperture opens, allowing surrounding liquid to enter the storage chamber. Fluid collection is initiated either by osmotic pressure [14] or by absorbing materials, such as hydrophilic fibers [15], a sponge [16], dehydrated hydrogel [17], or a collection bladder [18]. Some devices lack a closing mechanism, meaning the sampling process continues throughout the gastrointestinal tract [14]. However, most devices terminate the sampling process by closing the capsule’s aperture. This closure can be achieved by various mechanisms, including a bistable mechanism triggered by the expansion of an absorbing sponge [16], a spring-loaded latch that dissolves within 30 minutes, activating a piston that blocks the chamber inlet [19], a polydimethylsiloxane membrane that moves toward the aperture due to the expansion of dehydrated hydrogel [17], or a one-way valve that closes the aperture once the bladder is fully filled [18]. While the basic sampling processes have been briefly outlined in various articles [12, 13, 16], most of the capsules are not yet available commercially and none of the prototypes can currently be manufactured or reproduced, as the exact device specifications have not been fully detailed. Of all these capsules, only a few have been tested in humans [15, 18-20], in dogs [21], and, to a lesser extent, in pigs [14, 22], with limited success. This article presents a new capsule specifically designed for sampling the small intestine in pigs. The composition, specifications, and manufacturing process are thoroughly described, as is the protocol for administering the capsule to pigs, along with the methods used to validate the sampling location.

## II. Methods

### A. Capsule specifications

The sampling process can be divided in three phases. First, the capsule is to be administered orally, be swallowed, and transit through the stomach (phase 1). Second, it should open in the small intestine, collect a sample of the small intestine content, and close again (phase 2). Finally, it should remain sealed before being expelled with the feces (phase 3). As the capsule was intended to be first use in pigs, capsule’s requirements had to be specifically adapted to the specificity of pigs gastro intestinal anatomy. In addition, the capsule was intended to be used in young pigs (around 10 kg body weight), so the size and shape had to be adapted to the dimensions of the gastro-intestinal track at this age.

Unlike human, pig’s stomach has a very pronounced “C” shape, and the gastric cardia is very close to the pylorus [23]. There is also a transverse pyloric fold called the pyloric torus. This is located just before the pylorus and acts as a ‘gatekeeper’ to prevent large particles from entering the small intestine [24]. Due to these anatomical restrictions, pig’s gastric emptying is very slow, and only small amounts of stomach content leave the stomach at once [25]. The speed of gastric emptying is highly variable between pigs, and large solid particles (>1 cm) can remain in the stomach for several days [26]. Therefore, the capsule had to have a small diameter, ideally less than 10 mm, to optimise gastric passage (phase 1). At the same time, the capsule had to remain close while transiting through the stomach. As with other passive capsules, an enteric coating could keep the capsule sealed while passing through acidic conditions and open once the pH increases in the small intestine.

Utilizing commercially available enteric gelatin capsules could streamline development, so the sampling capsule should be designed to fit within commercial capsule sizes.

Once in the small intestine (phase 2), the capsule must be able to collect a sample. A passive collection system improves patient safety, allows for a larger sampling volume and simplifies the device. It also reduces the manufacturing cost and open the possibility for a single use device. After sampling, the device should close and remain sealed (phase 3) to avoid cross contamination while the capsule transits in the colon or remains in the feces. Closing the capsule must prevent any leakage of fluid and cross contamination until the device is evacuated naturally. As the purpose of the proposed capsule is to sample microbiota, the sample volume should be greater than 100µL. Sample should be easily collected by aspiration after opening of the capsule. To ensure the safety of the device, all materials in contact with the intestinal mucosa must be biocompatible and the shape of the capsule must not protrude at any stage of the process.

### B. Proposition of operating mode

Unlike humans, pigs do not voluntarily swallow solid capsules, so they must be delivered directly into the esophagus to avoid chewing [27]. For this, a dedicated esophageal sonde is designed for the capsule administration, and the contention method is adapted to not hamper the swallowing, i.e., a sling is used to restrain pigs. Then, taking into account the gastrointestinal characteristics of the pig and the above specifications, a 5-step operating mode is outlined. The capsule is initially encased by the upper part of an enteric coating, which keeps it sealed as long as the pH remains acidic (Fig. 1a). Upon reaching the small intestine, the gelatin-based upper section dissolves as the pH exceeds 6. This triggers the top of the capsule to open, allowing intestinal fluids to be drawn into the inner chamber by osmotic pressure (Fig. 1b). Once inside, the intestinal fluid at 38°C dissolves the gelatin glue securing a plunger in place (red arrows, Fig. 1c). The plugger then rapidly expands within seconds, pulling intestinal contents into the storage chamber (Fig. 1d). Finally, the capsule reseals as the plugger reaches the top of the capsule (Fig. 1e). The capsule is then excreted in the feces. A proper collection system is designed to facilitate capsule recovery in feces and reduce the risk of capsule destruction when pigs are housed in groups. To achieve this, the openings in the slatted floor were made narrower than the capsule’s diameter, and the overall slatted floor area was reduced [27].

**Fig. 1:**
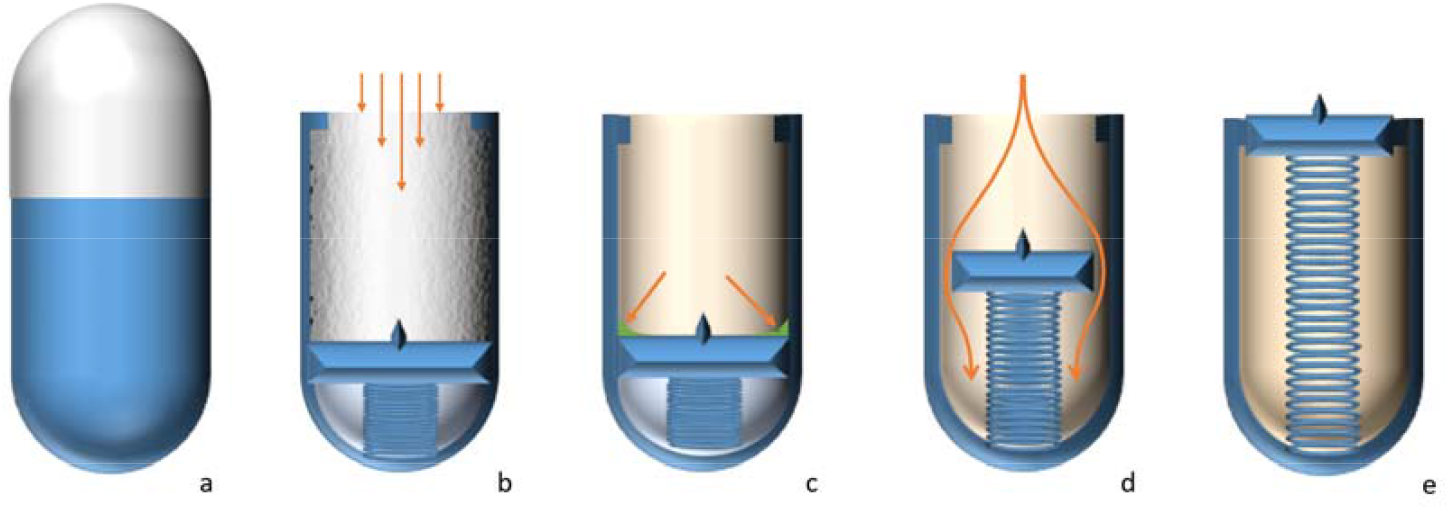
Mechanism of sampling. The capsule is initially covered by the upper part of an enteric capsule and remains sealed as long as the pH is acidic (1a). When the capsule reaches the small intestine, the gelatin-based top dissolves as the pH rises above 6. The top of the capsule opens and intestinal fluids are drawn into the inner chamber by the osmotic pressure (1b). The intestinal fluid at a temperature of 38°C dissolves the gelatin glue that was holding a plugger in the compression position (1c). As the plugger expands rapidly in a few seconds, it aspirates intestinal contents into the storage chamber (1d). The capsule closes again when the plug reaches the top of the capsule (1e).

### C. Key components of the capsule

The capsule is composed of four parts: a body, a plugger, a seal ring, and a spring (Fig. 2). The body is shaped like a hemi-capsule, measuring 20.0 mm in length, with an internal diameter of 6.4 mm and an external diameter of 7.2 mm. It features two internal narrowings, one at the top and one at the base, which ensure the storage chamber remains sealed before and after sample collection. The precise design of both the body and the plugger in stereolithographie (“.stl”) format are provided in Additional files 1 and 2, respectively. The plugger measures 6.1mm in length with a diameter of 3.6 mm. It includes a small button on top, which can be easily grasped with forceps to open the capsule and retrieve its contents after collection. A small notch on the plugger accommodates the O-ring (OR-1.50X2-NBR70, Rodi SA, Marly, Switzerland), which has an inner diameter of 1.5 mm and a thickness of 2 mm. The base of the plugger is designed to fit inside the spring (C-040.0440.0170.10-l, Dejex SA, Bienne, Switzerland), which measures 4.8 mm when compressed, 17 mm when released, and exerts a force of 0.44 N/mm. Before administration, the plugger compresses the spring and is held in place by a gelatin glue. The inner chamber of the body is filled with table salt, and the top is covered by an enteric capsule (Capsugel Enprotect® capsule size 0, Lonza, Colmar, France). After sampling, the released spring keeps the plugger at the top of the body, ensuring a watertight connection via the O-ring.

**Fig. 2:**
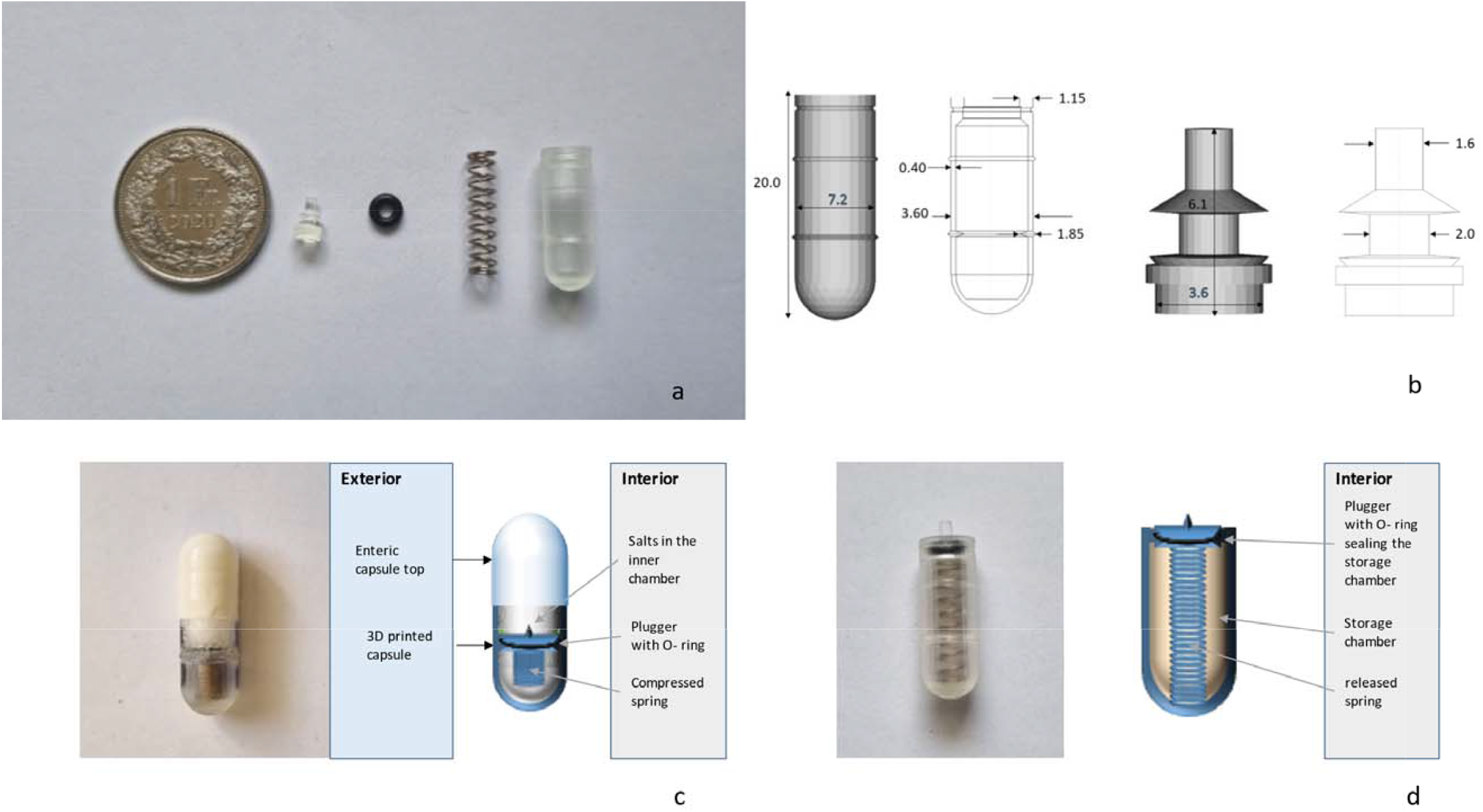
Components of the capsule. The four parts of the capsule (2a). Dimensions of the body and the plugger in mm (2b). Picture and scheme of the capsule, before administration (2c) and after retrieval (2d).

### D. Manufacturing

The capsules are produced through three main stages: printing, assembly, and finishing. First, the body and plugger are printed using a low-force stereolithography printer (Form 3, Formlabs, Berlin, Germany) with a biocompatible resin (surgical guide resin, Formlabs, Berlin, Germany) with a layer thickness of 0.1mm. Additional components, referred to as the “mushroom” due to its shape and the “claw” for its role in assembly, are also printed (Figure 3 a and b). The designs for these parts are available in stl format in Additional files 3 and 4, respectively. All printed parts are cleaned for 1 hour using Form Wash (Formlabs, Berlin, Germany) with 99% isopropanol. Afterward, a visual inspection is performed to ensure thorough cleaning, and the parts are left to dry for over 30 minutes. The parts are then post-cured at 60°C for 30 minutes with Form Cure (Formlabs, Berlin, Germany).

**Fig. 3:**
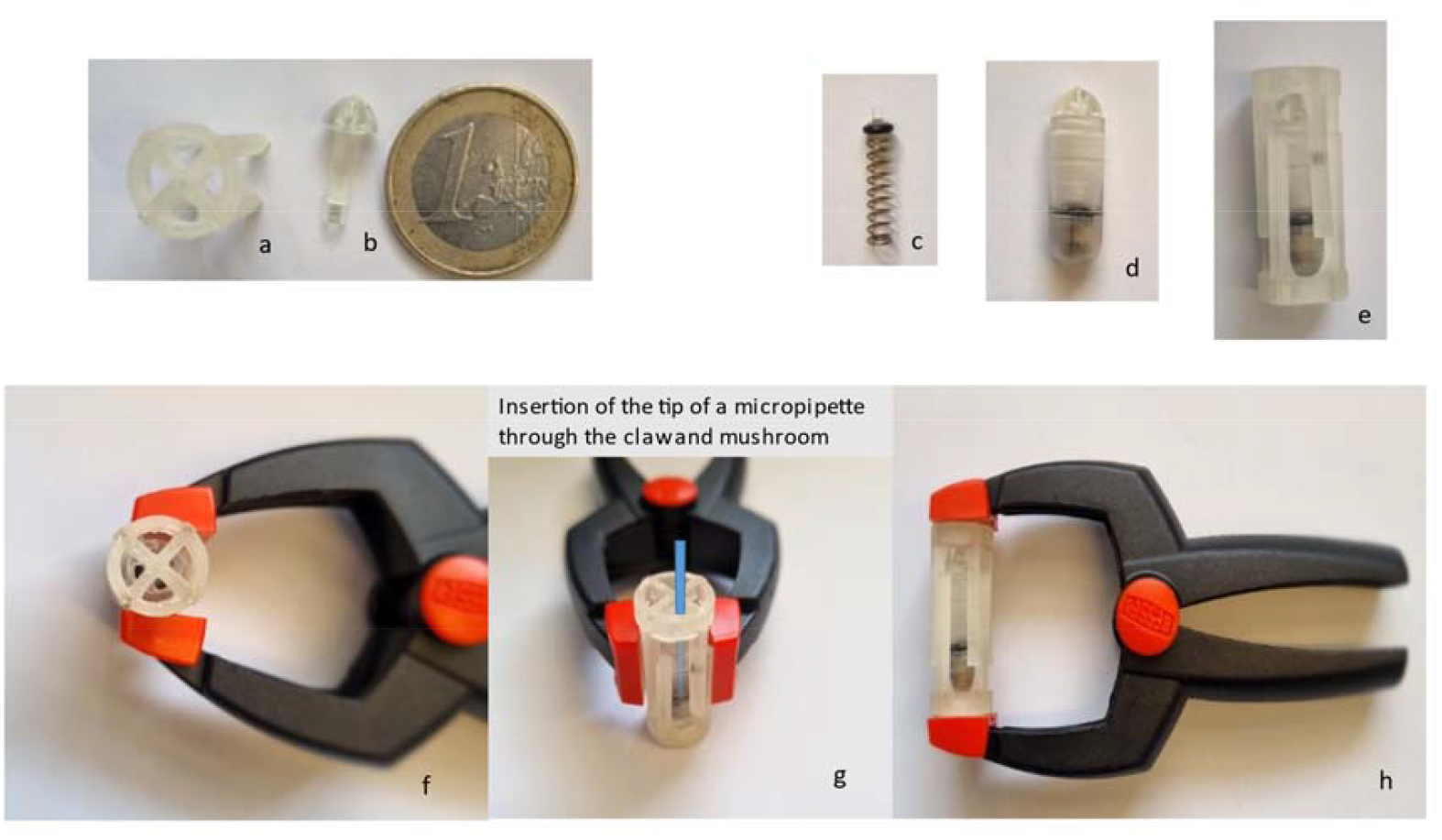
Assembling the capsule. It involves printing a mushroom (a) and two claws (b) to facilitate the process. The O-ring and spring are positioned on the plugger (c), which is then inserted into the body. The mushroom (d) and claws (e) hold the spring in a compressed state using a woodworking spring clip. Ethanol and gelatin glue are then injected directly onto the O-ring with a micropipette tip while the assembly is kept at 50°C in the oven. Finally, the assembly is dried for over 15 hours at 50°C (h).

During assembly, the O-ring is placed on the plugger, and a spring is inserted at its base (Figure 3c). The plugger assembly is then inserted into the body, with the spring being compressed and held in place by the mushroom and claws (Figure 3 d and e) using a woodworking spring clip (Figure 3f). The assembly is heated in an oven at 50°C. After 15 minutes in the oven, 20 µL of 70% ethanol (VWR chemicals, Dietikon, Switzerland) is injected at the base of the plugger using a micropipette (Figure 3g) to ensure even distribution of the gelatin glue around the plugger. The gelatin glue is prepared by dissolving 1.2 g of gelatin (Gelatin from porcine skin – gel strength 300, type A, Sigmaaldrich, Saint Louis, USA) in 6 mL of isotonic NaCl and melting it at 50°C for 45 minutes. Once the glue is liquid and homogenized, 80 µL is injected into the base of the plugger in the same manner as the ethanol. The assembly is then dried overnight in an oven at 50°C (Figure 3h) in vertical position.

After at least 15 hours, the claw and mushroom are gently removed, and the plugger remains compressed by the spring, held in place by the gelatin glue. The inner chamber is filled with table salt, and the enteric capsule top is fitted onto the body. Finally, the enteric top is sealed to the body by quickly dipping the base of the capsule into 70% ethanol and allowing it to dry. The capsule is then ready for administration.

### E. Experimental evaluations of the capsule

#### 1. In vitro validation

A first study assessed the sampling mechanism of the capsule *in vitro* under various simulated gastrointestinal environment as presented by [28]. In a nutshell, three *in vitro* assays were carried out to test the capsule in both low (pH≤4.2) and neutral pH (pH=7) solutions to simulate the gastric and the small intestine environment, respectively. In assay 1, the capsules (n=51) were placed in water at pH=2 for 30 min and then moved into water at pH=7 until all capsules had sampled an aliquot of the solution. In assay 2, the capsules (n=8) were left for 120 min at pH=2 and then moved to pH=7. In assay 3, the capsules (n=5) were kept for 2 h in a solution simulating gastric juices at pH=4.2 and then moved to a buffer at pH=7. In all three trials, none of the capsules started sampling under acidic conditions.

However, as per design, all capsules correctly collected samples under neutral conditions. The majority of the capsules (72.5 %, 100% and 100 %, in assay 1, 2 and 3, respectively) collected a sample within 60 minutes of being moved to the simulated small intestinal environment (pH=7). The sampling (opening and closing) took less than 10 seconds and sampled volumes ranging from 100 to 250 µL. The results of this first study were quite promising and revealed that the capsules performed well in all the simulated environments tested [28].

#### 2. In vivo validation

The capsules were initially tested in post-weaning pigs, as outlined by García Viñado et al. [29]. In summary, 26 pigs, each weighing either below 12 kg or between 12 and 20 kg, were administered two capsules and monitored over a period of three days. Administering swallowable devices to pigs poses significant challenges due to their anatomy, the need for dedicated materials for oesophagal intubation, and the requirement to restrain the animals without hampering the swallowing. Therefore, the protocol to administer the capsule followed García’s detailed procedures [27], which included administering a prokinetic agent (prucalopride) to promote gastric emptying, and using esophageal intubation, to address issues related to administration, gastric blockage, and capsule retrieval. After three days, the animals were euthanized for post-mortem sampling, allowing for direct collection of gut microbiota from five segments of GIT. This approach enabled a direct comparison between the microbial content of the gut and the samples collected by the capsules, and it therefore allowed for precise identification of the capsules’ sampling locations within the GIT. None of the pigs weighing less than 12kg excreted the capsules in the feces, and the majority of the capsules remained stuck in the stomach. Of the capsules administered to pigs above 12kg, 62.5% were successfully retrieved from the feces within 48 hours, and 46% were selected for microbiome analysis based on having a collected sample with a pH > 5.5. The bacterial composition from the capsules was compared to that of three segments of the small intestine, the large intestine, and the feces of the corresponding pigs. The bacterial composition within the capsules differed significantly from that of the large intestine and feces (P < 0.01), but it was not significantly different from that of the first segment of the small intestine (P > 0.10). This study provided the first evidence that the capsule effectively samples intestinal microbiota from the upper part of the small intestine in pigs weighing between 12 and 20kg.

In a similar study conducted with pigs weighing between 50 and 70 kg, results demonstrated that the capsules sampled the middle portion of the small intestine instead [27]. This difference in sampling location is likely due to variations in intestinal transit time. In growing pigs, the capsules may have opened and closed further along the intestine due to faster transit times. Indeed, Snoeck et al. (2004) observed that transit times were significantly prolonged immediately after weaning and returned to normal three weeks later [30].

## III. Discussion

This article provides comprehensive details on producing a capsule designed for sampling small intestine microbiota in pigs of various sizes. The capsule shares similarities with other passive sampling devices, as it uses an enteric coating that activates in response to the pH increase after passing through the stomach [14, 16-19, 22]. However, this capsule differs in that its sampling mechanism is triggered only after a gelatin glue dissolves, which delays the sampling process. On average, the mechanism is activated 30 to 60 minutes after the pH rises above 5.5. In addition, the sampling duration is very brief, lasting less than 10 seconds, ensuring that the sampling is confined to a specific location. However, as for the other passive sampling devices, the exact sampling location is not controlled, as it depends on the transit time through the gastrointestinal tract, which is influenced by the pig’s age, diet, and any underlying pathologies. This leads to both intra-individual and inter-individual variations. Nevertheless, the primary advantage of relying on a passive mechanism is that it provides a wide safety margin and lowers production costs, making it more suitable for a single-use device.

The capsule has also been validated in pigs as an effective tool for collecting small intestine content. Similar to other capsules [16, 18], we have shown that the microbiome extracted from the capsule sample is significantly different from that of faeces and colon, indicating that the capsule does not sample from these regions and there is no contamination after sampling. Additionally, we further confirmed the precise sampling location by directly comparing the microbiome of different sections of the small intestine with that collected by the capsule, showing that it samples the upper or middle segment depending on the size of the pig. It also demonstrates that the microbiome profile is not distorted by the time spent in the capsule during gastrointestinal transit. Therefore, the capsule serves as a virtual window into the small intestine, enabling the analysis of its contents while utilizing the natural pathway.

While this study focuses on the microbiome analysis of the capsule’s contents, the ultimate goal is to expand its application to analyze not only the microbiome but also the metabolome, viral activity, host proteome, and bile acid content within the intestines.

Additionally, the same sampling device could potentially be used to sample other sections of the intestine. The capsule is designed to fit any size 0 gelatin capsule, meaning that altering the properties of the hard gelatin top could allow sampling from the large intestine. Similarly, Shalon et al. (2023) [18] used different enteric coatings to sample three segments of the small intestine and the ascending colon in humans. Furthermore, the device can be tested in other species, revealing that exact sampling location may vary depending on the use case. Indeed, it could be valuable to compare the microbiome collected by the engineered Small Intestinal MicroBiome Aspiration (SIMBA™), which has already been safely administered to dogs [21], with that collected by the present capsule.

The production system may also require further optimization to streamline the process, enhance production efficiency, and enable scalability for larger batches. This could involve simplifying and automating the manufacturing process, which would significantly reduce manual labor and minimize the potential for human error. By automating key steps, the production cycle could become faster and more consistent, ultimately leading to higher quality and more reliable results.

## IV. Conclusions

One of the key advantages outlined in the article is the accessibility of the detailed production steps and the provided 3D printing file, which make it possible for other researchers and developers to easily suggest and test their own improvements. This transparency fosters an open-source approach to innovation, allowing users to modify and refine the design to better suit their needs or to push the technology further.

By making the manufacturing process openly available, the door is opened to a great deal of creativity and collaboration within the scientific community. Researchers and developers from various fields can build upon the existing work, leading to rapid advancements in both the design and function of the device. Ultimately, this collective effort could benefit everyone, making the device more accessible, cost-effective, and versatile for a wide range of applications.

## V. List of abbreviations

GIT: gastro intestinal tract
STL: stereolithography

## VI. Declarations

### A. Ethics approval

All experimental procedures were in compliance with Swiss animal welfare guidelines and were approved (No. 2021-39-FR, 2022-26-FR, 2022-39-FR) by the Cantonal Veterinary Office of Fribourg (Switzerland).

### B. Consent for publication

Not Applicable

### C. Availability of data and material

All data necessary to build a capsule are included in this published article (and its supplementary information files).

### F. Authors’ contributions

CO and BM conceived the prototype, CO, IGV and MT validated the procedure and performed the animal experiments, recorded the data, and collected and processed the capsules’ samples, and performed the statistical analysis. CO drafted the manuscript, and BM, MT, and IGV critically reviewed the manuscript. All authors read and approved the final manuscript.

